# gplas: a comprehensive tool for plasmid analysis using short-read graphs

**DOI:** 10.1101/835900

**Authors:** Sergio Arredondo-Alonso, Martin Bootsma, Yaïr Hein, Malbert R.C. Rogers, Jukka Corander, Rob JL Willems, Anita C. Schürch

## Abstract

**Summary:** Plasmids can horizontally transmit genetic traits, enabling rapid bacterial adaptation to new environments and hosts. Short-read whole-genome sequencing data is often applied to large-scale bacterial comparative genomics projects but the reconstruction of plasmids from these data is facing severe limitations, such as the inability to distinguish plasmids from each other in a bacterial genome. We developed gplas, a new approach to reliably separate plasmid contigs into discrete components using sequence composition, coverage, assembly graph information and clustering based on a pruned network of plasmid unitigs. Gplas facilitates the analysis of large numbers of bacterial isolates and allows a detailed analysis of plasmid epidemiology based solely on short read sequence data.

**Availability and implementation:** Gplas is written in R, Bash and uses a Snakemake pipeline as a workflow management system. Gplas is available under the GNU General Public License v3.0 at https://gitlab.com/sirarredondo/gplas.git

## 1 INTRODUCTION

A single bacterial cell can harbor several distinct plasmids, however, current plasmid prediction tools from short read WGS often have a binary outcome (plasmid or chromosome). To bin predicted plasmids into discrete entities, we built a new method based on the following concepts: i) contigs of the same plasmid have a uniform sequence coverage ^1,10^, ii) plasmid paths in the assembly graph can be searched for using a greedy approach^8^ and iii) removal of repeat units from the plasmid graphs disconnects the graph into independent components^12^.

Here, we refined these ideas and introduce the concept of unitigs co-occurrence to create a pruned plasmidome network. Using an unsupervised approach, the network is queried to find highly connected nodes corresponding to sequences belonging to the same discrete plasmid unit, representing a single plasmid. We show that our approach outperforms other de-novo and reference-based tools and fully automates the reconstruction of plasmids.

## 2 MATERIALS AND METHODS

### 2.1 Gplas algorithm

Given a short-read assembly graph (gfa format), segments (nodes) and edges (links) are extracted from the graph. Gplas uses mlplasmids (version 1.0.0, prediction threshold = 0.5) or plasflow (version 1.1, prediction threshold = 0.7) to classify segments as plasmid- or chromosome-derived and selects segments with an in- and out-degree of 1 (unitigs) ^2, 7^. The k-mer coverage standard deviation (k-mer sd) of the chromosome-derived unitigs is computed to quantify the fluctuation in the coverage of segments belonging to the same replicon unit. Plasmid-derived unitigs are considered to search for plasmid walks with a similar coverage and composition using a greedy approach (Supplementary Methods). Gplas creates a plasmidome network (undirected graph) in which nodes correspond to plasmid unitigs and edges are drawn based on the co-existence of the nodes in the solution space using R packages igraph and^4^ ggraph(https://github.com/thomasp85/ggraph.git). Markov clustering algorithm is used to query the plasmidome network and retrieve clusters corresponding to discrete plasmid units in an unsupervised fashion^11^. The output consists of plasmid contigs binned into distinct components, representing the different plasmids present in the bacterial isolate. Complete description of the algorithm is available in Supplementary Methods.

### 2.2 Benchmarking dataset

Gplas was benchmarked against current existing tools to bin plasmid contigs from short-read WGS: i) plasmidSPAdes (de-novo based approach, version 3.12)^1^, ii) mob-recon (reference-based approach, version 1.4.9.1)^9^ and iii) hyasp (hybrid approach, version 1.0.0)^8^. To evaluate the binning tools, we selected a set of 28 genomes with short- and long-read WGS available including 106 plasmids from 10 different bacterial species which were not present in the databases or training sets of the tools (Supplementary Methods, Supplementary Table S1)^3, 5, 6, 13^.

For each component reported by gplas, we can consider *n* as the total number of nodes present in the component. Then, we can define *C* as:

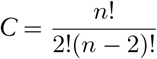

*C* corresponds to the total number of pair-pair connections between nodes of a particular component. We consider as true positive connections (*TPC*), pair-pair connections linking to nodes belonging to the same replicon sequence in contrast to false positive connections (*FPC*) in which connections link to nodes from different replicon sequences. Let *n_pc_* be the total number of nodes from the most predominant replicon sequence present in the component and *n*_*rep*_ the total number of nodes forming that replicon sequence. We then define two metrics commonly used in metagenomics for binning evaluation: i) precision and completeness (Supplementary Methods).

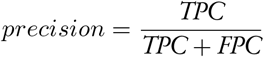

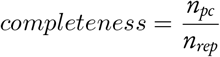

## 3 RESULTS

Gplas in combination with mlplasmids obtained the highest average precision (0.85) indicating that the predicted components were mostly formed by nodes belonging to the same discrete plasmid unit (Table 1 and Figure S1). The reported average completeness value (0.73) showed that most of the nodes from a single plasmid were recovered as a discrete plasmid component by gplas (Table 1 and Figure S2). We observed a decline in the performance of gplas in combination with mlplasmids (precision = 0.71, completeness = 0.68) when considering uniquely complex components (> 1 connection) which indicated merging problems of large plasmids with a similar k-mer coverage (Figure S3, Supplementary Results). However, in all cases the performance of gplas in combination with mlplasmids performed better than other de-novo and reference-based tools tested here (Table 1). To show the potential of gplas in combination with mlplasmids, we showcase the performance of our approach in two distinct bacterial isolates (Supplementary Results).

**Table 1.** Gplas benchmarking. *Components > 1 connection

| Tool | Precision | Completeness |
| --- | --- | --- |
| gplas - mlplasmids | 0.85/0.71* | 0.73/0.68* |
| gplas - plasflow | 0.63/0.45* | 0.55/0.42* |
| hyasp | 0.53/0.34* | 0.42/0.34* |
| mob-recon | 0.62/0.61* | 0.50/0.56* |
| plasmidSPAdes | 0.47/0.20* | 0.71/0.67* |

Mlplasmids only contains a limited range of species models (Supplementary Methods). For other bacterial species, we observed that plasflow probabilities in combination with gplas performed better than the other de-novo approaches but also introduced bias when wrongly predicting chromosome contigs as plasmid nodes (Table 1 and Figure S1), thereby creating chromosome and plasmid chimeras (precision = 0.63).

## 4 DISCUSSION

We present a new tool called gplas which enables the binning and a detailed analysis workflow of binary classified plasmid contigs into discrete plasmid units by relying on the structure of the assembly graph, k-mer information and clustering of a pruned plasmidome network. A limitation of the presented approach is the generation of chimaeras resulting from plasmids with similar k-mer profiles and sequence coverage and sharing a repeat unit, such as a transposase or an IS element. These cases cannot be unambiguously solved. Here, we integrated and extended upon features to predict plasmid sequences and exploit the information present in short-read graphs to automate the reconstruction of plasmids.

## ACKNOWLEDGEMENTS

We would like to thank Dr. Bryan Wee for testing and contributing to the development of gplas.

## REFERENCES

1. Antipov, D. et al. (2016). plasmidSPAdes: Assembling plasmids from whole genome sequencing data. Bioinformatics, 32, 3380–3387.

2. Arredondo-Alonso, S. et al. (2018). mlplasmids: a user-friendly tool to predict plasmid- and chromosome-derived sequences for single species. Microb Genom.

3. Arredondo-Alonso, S. et al. (2019). Genomes of a major nosocomial pathogen enterococcus faecium are shaped by adaptive evolution of the chromosome and plasmidome. bioRxiv.

4. Csardi, G. et al. (2006). The igraph software package for complex network research. InterJournal, Complex Systems, 1695(5), 1–9.

5. De Maio, N. et al. (2019). Comparison of long-read sequencing technologies in the hybrid assembly of complex bacterial genomes.

6. Decano, A. G. et al. (2019). Complete assembly of escherichia coli sequence type 131 genomes using long reads demonstrates antibiotic resistance gene variation within diverse plasmid and chromosomal contexts. mSphere, 4(3).

7. Krawczyk, P. S. et al. (2018). PlasFlow: predicting plasmid sequences in metagenomic data using genome signatures. Nucleic Acids Res., 46(6), e35.

8. Müller, R. and Chauve, C. (2019). HyAsP, a greedy tool for plasmids identification.

9. Robertson, J. and Nash, J. H. E. (2018). MOB-suite: software tools for clustering, reconstruction and typing of plasmids from draft assemblies. Microb Genom, 4(8).

10. Rozov, R. et al. (2015). Recycler: an algorithm for detecting plasmids from de novo assembly graphs. BioRxiv, page 029926.

11. Van Dongen, S. (2008). Graph clustering via a discrete uncoupling process. SIAM J. Matrix Anal. Appl., 30(1), 121–141.

12. Vielva, L. et al. (2017). PLACNETw: a web-based tool for plasmid reconstruction from bacterial genomes. Bioinformatics.

13. Wick, R. R. et al. (2017). Completing bacterial genome assemblies with multiplex MinION sequencing. Microbial Genomics, 3(10).

