## Supplementary Materials for "gplas: a comprehensive tool for plasmid analysis using short-read graphs"

### Supplementary material from gplas: a comprehensive tool for plasmid analysis using short-read graphs

#### SUPPLEMENTARY METHODS S1

Gplas requires a single input corresponding to a graph in gfa format (version 1.0) (https://github.com/GFA-spec/GFA-spec). We can define the nodes $N$ present in a graph $G$ by:

$$N = \{N_1, N_2, ..., N_n\},$$

i.e., the number of nodes in $G$, $|N|$ equals $n$. Nodes can also be referred to as segments, contigs or vertices. Furthermore, we can define the set of links $L$ as the set of directed connections between two elements of $N$:

$$L = \{(i, j) : |1 \leq i, j \leq n \quad \text{with a link from } N_i \text{ to } N_j\}.$$

Links can also be referred as edges. We also denote the edge from vertex $v$ to vertex $w$ by $e(v, w)$. We can define the graph $G$ given by the user as:

$$G = (N, L).$$

For each $v \in N$, we define the in-degree $d^i(v)$ and the out-degree $d^o(v)$ as the number of edges in $L$ towards and from vertex $v$ respectively. We define $|v|$ as the length of segment $v$, i.e., the number of nucleotides of segment $v$.

Mlplasmids (version 1.0.0) provides for each $v \in N$ a probability whether $v$ is plasmid-derived if the given $G$ corresponds to a bacterial species included in the tool (*Enterococcus faecium*, *Klebsiella pneumoniae* or *Escherichia coli*). To generalize the prediction to other bacterial species, the probabilily that the segment is plasmid-derived can also be derived using plasflow (version 1.1) which provides a metagenomics classifier to retrieve plasmid-derived sequences. For either method, we define $\pi(v)$ as the probability that segment $v$ is plasmid-derived.

For a graph $G$ generated by SPAdes, the k-mer count information of node $N_i$ is extracted from the "kc" tag present in the header line of $N_i$ for each node $1 \leq i \leq n$ . The k-mer count is divided by $|N_i|$ and normalised against the median k-mer count. If $G$ was generated from Unicycler, the normalised depth of $N_i$ (k-mer coverage) is retrieved from the tag "dp" which is already present in the header line of $N_i$. In both cases, this value is further considered and defined as the *coverage* of a particular node $v \in N$, and denoted by $c(v)$.

Let $t$ be the threshold for posterior probability of the plasmid class $\pi$ reported by mlplasmids or plasflow. We then define $\mathcal{P}$ as the set of plasmid-predicted contigs by:

$$\mathcal{P} = \left\{v \in N : \pi(v) \geq t, d^i(v) = d^o(v) = 1, |v| \geq 1\, kbp\right\}.$$

And $\mathcal{C}$ as the set of chromosome-predicted contigs by:

$$\mathcal{C} = \left\{v \in N : \pi(v) < t, d^i(v) = d^o(v) = 1, |v| \geq 1\, kbp\right\}.$$

$|\mathcal{P}|$ and $|\mathcal{C}|$ are the number of contigs that are plasmid-predicted and chromosome-predicted, respectively. For the set of chromosome-predicted contigs, we define the average coverage $\mu_\mathcal{C}$ and the variance of the coverage $s^2_\mathcal{C}$ by:

$$\mu_\mathcal{C} = \frac{1}{|\mathcal{C}|}\sum_{v\in\mathcal{C}} c(v)$$

$$s^2_\mathcal{C} = \frac{1}{|\mathcal{C}|-1}\sum_{v\in\mathcal{C}}(c(v)-\mu_\mathcal{C})^2.$$

For a graph $G = (N, L)$, a walk of length $k \in \mathbb{N}$ is defined as a finite sequence of alternating vertices and edges $(v_0, e(v_0, v_1), v_1, e(v_1, v_2), \ldots, e(v_{k-1}v_k), v_k)$ with $v_i \in N$ for $0 \leq i \leq k$ and edge $e(v_{i-1}v_i)$ is the edge in $L$ from $v_{i-1}$ to $v_i$ for all $1 \leq i \leq k$. We denote the set of vertices $\{v_0, v_1, \ldots, v_k\}$ of a walk $W$ by $V(W)$.

Here we consider a special class of walks with the following properties:

1. $v_0 \in \mathcal{P}$, i.e., the walk starts from a segment which is predicted to be plasmid-derived.
2. $k \leq M = 100$, i.e, we only consider of walks of length less or equal to $M = 100$.
3. $v_i \neq v_0$ for $1 \leq i < k$ i.e., the walks either does not return to the starting vertex or ends when it returns to the starting vertex.

If we have a walk $W = (v_0, e(v_0, v_1), v_1, e(v_1, v_2), \ldots, e(v_{k-1}, v_k), v_k)$ which can be extended, i.e., $k < M$ and $v_k \neq v_0$, we want to quantify whether a node $v$ for which the edge $e(v_k, v)$ exists, is a likely extension of the walk or not. We call such a node $v$ a candidate extension node. Slightly abusing notation, we define the average coverage of walk $W$, $c(W)$, as:

$$c(W) = \frac{1}{|V(W)\bigcap(\mathcal{C}\cup\mathcal{P})|}\sum_{v\in(V(W)\bigcap(\mathcal{C}\cup\mathcal{P}))} c(v),$$

i.e., we base the average coverage of $W$ only on the coverage of the nodes of $W$ which are at least 1 *kbp* long and have an in-degree and out-degree equal to one.

Alternatively, we can also determine the average coverage by weighting each contig by its length. We then obtain an alternative average coverage of walk $W$, denoted by $\tilde{c}(W)$, defined by:

$$\tilde{c}(W) = \frac{\sum\limits_{v\in(V(W)\bigcap(\mathcal{C}\cup\mathcal{P}))} c(v)|v|}{\sum\limits_{v\in(V(W)\bigcap(\mathcal{C}\cup\mathcal{P}))} |v|}.$$

If the candidate extension node $v$ belongs to either $\mathcal{C}$ or $\mathcal{P}$, we want to determine how similar $c(v)$ and $c(W)$ are. For simplicity, we assume that the variance in the coverage of a node due to chance events related to the sequencing process is $s^2_\mathcal{C}$, i.e., the variance in the coverage of the chromosome-predicted contigs, more precisely, we assume that the coverage of the next contig of the walk is distributed according to a normal distribution with mean $c(W)$ and variance $s^2_\mathcal{C}$ if the walk and the candidate extension node belong to the same plasmid. We define the similarity in coverage between the walk $W$ and the candidate extension node $v$, $S(W, v)$ as:

$$S(W,v) := \Phi(\frac{c(v)-c(w)}{s_\mathcal{C}}+1) - \Phi(\frac{c(v)-c(w)}{s_\mathcal{C}}-1)$$

with $\Phi$ the cumulative distribution function of the standard normal distribution. This means that the closer $c(v)$ is to $c(W)$, the higher $S(W, v)$.

The similarity $S(W, v)$ is based only on the *coverage*. In order to avoid chimeras between chromosomal and large plasmid replicons with a similar *coverage*, we define a score $g(W, v)$ of each candidate extension node by:

$$g(W, v) = \begin{cases} 0.25 & \text{if } v \notin (\mathcal{P} \cup \mathcal{C}) \\ S(W, v) \cdot \pi(v) & \text{if } v \in (\mathcal{P} \cup \mathcal{C}) \end{cases}$$

Candidate extension nodes $v$ assigned with a score $g$ of 0.25 may belong to repeat units such as transposases or IS elements and thus $\pi(v)$ and $S(W, v)$ cannot be confidently estimated. Additionally, we apply a filtering threshold $\xi$ (default $\xi = 0.10$) to avoid the selection of edges potentially leading to the creation of replicon chimeras. This threshold can be tuned by the user to accept or reject a higher number of connections.

Let $E(W)$ be the set of candidate extension nodes of walk $W$. We define the function $h_W(v)$ by:

$$h_W(v) = \begin{cases} 1 & \text{if } g(W, v) \geq \xi \\ 0 & \text{if } g(W, v) < \xi \end{cases},$$

i.e., $h_W(v)$ equals one, if the candidate extension node has a score higher or equal to the threshold $\xi$.

If $h_W(v) = 0$ for all $v \in E(W)$, we have reached a dead-end, otherwise we select the extension $v \in E(W)$ with probability

$$\frac{g(W, v) h_W(v)}{\sum_{v' \in E} g(W, v') h_W(v')}$$

This results in the walk $W'$ which is the walk $W$ extended with the node $v$, i.e., $W' = (v_0, e(v_0, v_1), v_1, e(v_1, v_2), \ldots, e(v_{k-1}, v_k),$ Starting from a node which is plasmid-predicted, we repeat this extension procedure of the walk until we reach one of the following scenarios:

1. There are no outgoing links from the last element of $W$ i.e., we have reached a dead-end.
2. The last node incorporated in $W$ corresponds to the starting node of the walk.
3. The length of the walk $|W|$ exceeds $M = 100$.

For each $v \in \mathcal{P}$, we generate $K$ walks (default $K = 20$, but tunable by the user).

We use these $|\mathcal{P}|K$ walks, denoted by $\{W_1, W_2, \ldots W_{|\mathcal{P}|K}\}$ to generate a new pruned plasmidome network $G_{\mathcal{P}}(N_{\mathcal{P}}, L_{\mathcal{P}})$. The nodes of $G_{\mathcal{P}}$, denoted by $N_{\mathcal{P}}$ are all nodes which are present in at least one walk, i.e., $N_{\mathcal{P}} = \bigcup_{i=1}^{|\mathcal{P}|K} V(W_i)$.

To define the edges of $G_P$, denoted by $L_{\mathcal{P}}$, we first define for two nodes $v, w \in \mathcal{P} \cup \mathcal{C}$, a set $J(v, w)$ which describes in which walks both $v$ and $w$ are present, i.e.,

$$J(v, w) := \{i \in \mathbb{N} : 1 \leq i \leq |\mathcal{P}|K, v \in V(W_i) \text{ and } w \in V(W_i)\}.$$

Let $|J(v, w)|$ be the number of walks in which both $v$ and $w$ are present. We define a positive parameter $x \in \mathbb{R}$, with default value $x = 0.1$, which specifies the minimum number of walks in which two nodes $v$ and $w$ are both present in order to let the edge $e(v, w)$ be an element of $L_{\mathcal{P}}$. An edge $e(v, w)$ belong to $L_{\mathcal{P}}$ if $|J(v, w)| \geq xK$, i.e.,

$$L_{\mathcal{P}} = \{e(v, w) : v, w \in \mathcal{P} \cup \mathcal{C} \text{ and } |J(v, w)| \geq xK\}.$$

Note that the graph $G_{\mathcal{P}}(N_{\mathcal{P}}, L_{\mathcal{P}})$ is an undirected graph, as whenever $|J(v, w)| \geq x$, then $|J(w, v)| \geq x$ as well.

With this approach we create non-directed links between plasmid unitigs avoiding intermediary links connecting to nodes not present in $\mathcal{P} \cup \mathcal{C}$. These intermediary nodes can correspond to transposases or repetitive elements that are shared between replicon sequences in $G$.

The plasmidome network is considered as an undirected graph and represented in the R package igraph (version 1.2.4.1) (Csardi et al., 2006) and visualized using R package ggraph (version 1.0.2, https://github.com/thomasp85/ggraph.git).

Finally, $G_{\mathcal{P}}$ is queried using Markov clustering algorithm (mcl) available in the R package mcl (addLoops = FALSE, allow1 = TRUE) (version 1.0) to retrieve highly-connected nodes present in the plasmidome network (Van Dongen, 2008). Mcl finds cluster structure graph by simulating random walks considering two matrix operators, expansion and inflation. These two operators transform the set of probabilities of visiting nodes in $G_{\mathcal{P}}$. Inflation modifies the probabilities associated to the set of random walks of connecting one to a particular node and by this favouring more likely walks. Expansion increases the length of the random walks. Since nodes belonging to the same cluster in $G_{\mathcal{P}}$ share several nodes, the probabilities of nodes belonging to the same cluster will be higher since there are several ways of traversing nodes of the same cluster. In short, random walks in $G_{\mathcal{P}}$ will infrequently traverse natural clusters.

Nodes from $G_{\mathcal{P}}$ from a specific cluster reported by mcl are considered and reported as belonging to the same discrete plasmid sequence.

**SUPPLEMENTARY METHODS S2**

**Data:** Graph $G$

**Result:** Co-occurrence plasmidome network $G_{\mathcal{P}}$. Assignation of plasmid nodes $N_{\mathcal{P}}$ into different components

**Initialization**;

Extract nodes $N$ and links $L$ from $G$;

Divide $N$ as collection of plasmid-derived nodes $\mathcal{P}$ and chromosome-derived nodes $\mathcal{C}$ using mlplasmids or plasflow;

Discard $\mathcal{P}$ and $\mathcal{C}$ with an $d^i(v)$ and $d^o(v)$ != 1 and length < 1 kbp;

Determine the $s^2_{\mathcal{C}}$ of $\mathcal{C}$ based on the k-mer coverage;

```
foreach v0 ∈ P do
    Search through all the possible plasmid-like walks W starting from v0;
    for W in number of walks do
        while ∃ eligible extension E(W) do
            Consider the last v in W
            Retrieve all candidate extensions E(W)
            Compute gplas scores g(W,v) of E(W)
            Filter E(W) with a g(W,v) < ξ (default = 0.1, tunable by the user)
            Sample a E(W) based on the vector g(W,v)
            Extension of W using the selected v
        end
        Create a new set of links L_P connecting N_P in W;
        Reinitialize W considering again v0 as first element;
    end
end
Filter out L_P with a frequency < x (default = 0.1, tunable by the user) ;
Create a novel G_P using N_P and L_P;
Use markov-clustering algorithm MCL to query G_P;
Assignation of N_P in different components based on MCL output;
```

**Algorithm 1:** Gplas pseudocode

**SUPPLEMENTARY METHODS S3**

*Benchmarking Dataset*

To evaluate the performance of gplas against existing plasmid binning tools, we selected a set of 28 genomes with short- and long-read WGS available including 106 plasmids from 10 different bacterial species (Supplementary Table S1)[2–4, 11]. These genomes were selected due to their release date (after June 2017) to avoid any bias in favour of reference-based approaches as a result of including plasmids present in the databases of the tools. Importantly, these genomes were not part of the training sets of mlplasmids or plasflow.

We trimmed short-reads using trim galore (version 0.6.1) and determined a minimum quality phred score of 20[6]. Long-reads were filtered out using filtlong (v0.2.0) (https://github.com/rrwick/Filtlong.git) specifying a minimum length of 1 kbp, removing 10% of the worst base reads, filtering out reads with a mean quality weight inferior to 20 using short-reads as references and finally keeping a number of kbp corresponding to a genome coverage of 20x. We subsequently used Unicycler (version v0.4.7) using SPAdes (version 3.12.0) to perform a hybrid assembly and obtain complete genomes[12]. If the assembly resulted in an non-complete genome, we retrieved the uncompleted path using Bandage (version 0.8.1)[10] and used filtlong (v0.2.0) indicating the non-completed path as external reference and selecting reads from that path until reaching a path coverage of 10x. Previous filtered long-reads and reads corresponding to the non-completed path were merged and passed to Unicycler to rerun the hybrid assembly and obtain a complete genome.

**SUPPLEMENTARY METHODS S4**

*Benchmarking tools*

PlasmidSPAdes (from SPAdes 3.12) was run using default parameters and specifying the trimmed short-reads [1]. Hyasp (version 1.0.0) was run using the flag "–bin" and specifying as input the graph created by Unicycler using only short-reads and after removing overlaps (002_overlaps_removed.gfa) [7]. We created the database proposed by hyasp authors to identify plasmid seeds using the file "ncbi_database_genes.fasta" provided in the hyasp github repo (https://github.com/cchauve/HyAsP). Mob-recon (version 1.4.9.1) was run using default values with the proposed database (https://github.com/phac-nml/mob-suite) after first-installation of mob-suite [8]. Nodes from the short-read graph (002_overlaps_removed.gfa) were used as input for mob-recon. Gplas (version 0.4.0) was run with default values (f = 0.1, x = 20, mlplasmids threshold = 0.5, plasflow threshold = 0.7) indicating the short-read graph as input (002_overlaps_removed.gfa).

We used Quast (version 4.6.3) to map nodes predicted as belonging to the same component against the complete genomes from the same bacterial isolate [5]. To determine the precision and completeness (see section below) of the predictions, we only considered nodes with a length larger than 1 kbp, mapping unambiguously to a replicon sequence and thus excluding transposases and other repetitive sequences mapping totally or partially to more than one genome sequence.

**SUPPLEMENTARY METHODS S5**

*Benchmarking metrics*

The purity of the predicted components (precision) was assessed using the number of connections from a discrete genomic unit present in the component. The choice of connections rather than contigs as metric unit was selected on the basis of reporting the presence of more than one discrete genome, either chromosome or plasmid, merged into the same component.

For each component, we determined which reference replicon (either plasmid or chromosome) had a larger representation (in number of contigs). Completeness of the component was calculated as the fraction of the contigs recovered from the most predominant reference replicon. Components in which the most predominant reference replicon was the chromosome unit were assigned with a precision and completeness of 0. We strongly penalized these components predictions since they are mostly formed and contaminated by chromosome-derived sequences.

Some of the predicted components were small plasmids with a single connection corresponding to a self-loop and mostly resulted in precision and completeness of values of 1.0. These components can mask the problems of binning present in medium and large plasmids formed by several plasmid unitigs and with a similar k-mer coverage. To elucidate the performance of the tools in these cases, we also reported the average precision and completeness of the tools in predicted components uniquely with more than one connection.

**SUPPLEMENTARY METHODS S6**

*Gplas output files*

- **results/*results.tab**: Tab delimited file containing the full output information retrieved by gplas. The file contains the following columns: contig number, probability of being chromosome-derived, probability of being plasmid-derived, class prediction, contig name, k-mer coverage, length, component assigned.
- **results/*components.tab**: Tab delimited file containing the bin prediction reported by gplas with the following columns: contig number, component assignation
- **results/*component*.fasta**: Fasta file generated per each component containing the nodes assigned to them.
- **results/*plasmidome_network.png**: Png file of the plasmidome network generated by gplas after cre-

ating an undirected graph considering as nodes the plasmid unitigs and as edges the co-occurrence of these nodes in the space of solutions.

- **walks/*solutions.csv**: Comma-separated file containing the plasmid-like walks generated by gplas.
- **walks/*connections.tab**: Tab delimited file containing the following information: Variance factor, number of tries selected by the user, try number, elongation number, plasmid starting node, last node of the walk, possible outcoming node, probplasmid, probcoverage, gplas score, frequency of the score, selection or discardance of the connection. This file may facilitate the visualization of the walks generated by gplas as exemplified in Figures S8 and S9.

#### SUPPLEMENTARY RESULTS S1

*Gplas showcase: K. pneumoniae KSB1_7G*

To show the potential of gplas to bin contigs into discrete plasmid units, we showcase the isolate *Klebsiella pneumoniae* KSB1_7G (Supplementary Table S1). This genome has several features that allows to showcase the features used by gplas (Table S2): i) presence of two discrete components showing a similar k-mer coverage and containing plasmid starting nodes, ii) a low number of nodes that reduces the total number of walks, indispensable for visualization purposes and iii) no dead-ends present in the short-read graph.

| KSB1_7G short-read graph | Class | Nodes | Edges | Dead-ends |
|---|---|---|---|---|
| Component A | Chromosome | 74 | 100 | 0 |
| Component B | Plasmid 1 | 9 | 12 | 0 |
| Component C | Plasmid 2 | 1 | 1 | 0 |

**Table S2.** KSB1_7G summary short-read graph stats

Component B (Figure S5, left component) and C (Figure S5, right component) correspond to two discrete plasmid sequences with an approximated length of 161 kbp and 112 kbp respectively (Figure S5). Component C has a unique edge corresponding to a self-loop since that plasmid sequence has circularization signatures present in the boundaries of the node 15.

Gplas uses mlplasmids (or plasflow, in case a bacterial species not listed in mlplasmids is chosen) to predict plasmid-derived and chromosome-derived nodes. Predicted plasmid nodes with an in- out-degree equal to 1 (unitigs) are considered as plasmid starting nodes. Chromosome-derived nodes corresponding to unitigs are used to calculate the k-mer coverage standard deviation (0.034) present in nodes belonging to the same replicon. Plasmid starting-nodes (15, 18, 20, 24, 25, 26, 27) are considered to search for plasmid-like walks.

For the purpose of visualization procedures, we create a space search of 5 solutions per each plasmid starting node. We use the plasmid starting node 18+ to show the workflow of gplas:

- 18+ is the first node of the walk.
- The coverage of the walk corresponds to 1.35.
- The unique outcoming edge (18+,33-) corresponds to a connection with a repeat unit since 33- has two incoming and two outcoming edges (in-degree and out-degree of 2) resulting in a coverage of 2.70. This outcoming edge (18+,33-) is assigned with a default gplas score of 0.25 corresponding to a default k-mer composition and coverage score of 0.5 respectively.
- We sample the vector of gplas scores to select for an outcoming edge to elongate the walk. In this case, only a single edge can be selected to elongate the walk.

- We include the node (33-) from the outcoming edge (18+,33-). The first elongation results in the walk: 18+,33-.
- We update the k-mer coverage of the walk. In this case, the coverage of the walk remains 1.35 since the k-mer coverage of 33- is not considered. Only unitig nodes with a length larger than 1 kbp are considered in this step.
- There are two outcoming edges from the node 33- : i) 33-, 20- and ii) 33-, 24+.
- We calculate the probcoverage and retrieve the probplasmid of the outcoming edges. In this case, the edges connect to nodes showing a similar probcoverage (Figure S6 and S7) since both are part of the same replicon. We retrieve the probplasmid corresponding to the probabilities of being plasmid-derived: 0.86 (20-) and 0.94 (24+). This results in a gplas score of 0.31 (33-, 20-) and 0.48 (33-,24+).
- We sample one of the outcoming two edges and elongate the walk with the selected connection.
- In this case, we update the k-mer coverage of the walk using 18- and 20- or 24+, depending on which outcoming node was selected.

We follow this procedure until reaching the following three scenarios: i) all outcoming edges have a final gplas score lower than the filtering threshold (default = 0.10), ii) no outcoming edges are available to elongate the walk (e.g. dead-end) or iii) the last node incorporated in the walk corresponds to the initial plasmid starting node (circularization signature). We repeat this procedure 5 times (only for visualization purposes) and end up with the solutions illustrated in Figure S8. We find the solution: 18+,33-,20-,31-,25-,31+,18+ repeated in the 5 panels.

Interestingly, we find the same set of solutions if the plasmid search is initialized from the opposite direction 18- (Figure S9).

We repeat the same approach with the other plasmid starting nodes (15, 20, 24, 25, 26 and 27) to obtain our space of solutions.

We create a new plasmidome network consisting of an undirected graph in which nodes correspond to the plasmid starting nodes and edges to co-occurrence of these nodes in the solutions found by gplas (Figure S10). Using markov clustering algorithm[9], we search for highly-connected plasmid starting nodes in the plasmidome network using an unsupervised approach. Nodes categorized as belonging to the same cluster are reported by gplas as belonging to the same discrete plasmid sequence (Table S3).

In Figure S10, we can observe two different components present in the plasmidome network. Nodes belonging to these components are binned using markov clustering and reported in Table S3.

| Plasmid starting node | Component |
|---|---|
| 15 | 2 |
| 18 | 1 |
| 20 | 1 |
| 24 | 1 |
| 25 | 1 |
| 26 | 1 |
| 27 | 1 |

**Table S3.** Assignation of plasmid starting nodes into different components (bins).

In this simplified example in which we only requested for the search of 5 plasmid-like walks per starting node, two different components were obtained. Component 2 contains a single node, 15, with a self-loop indicating the presence of circularization signatures (precision 1.0, completeness = 1.0). Interestingly, even though node 15 has a k-mer coverage similar to other plasmid starting nodes, the structure of the original assembly graph indicated that it corresponded to an independent plasmid sequence. This highlights the importance of searching for walks using the original graph rather than simply binning nodes based on similar k-mer coverage or composition. Component 1 (precision 1.0, completeness = 1.0) belongs to plasmid 2 (Table S3).

#### SUPPLEMENTARY RESULTS S2

*Gplas showcase on plasmids with a similar k-mer coverage*

To highlight the main limitation of gplas, we show the results obtained for the *K. pneumoniae* isolate SAMN10819819 (Supplementary Table S1). This isolate contained five plasmids (Figure S11), and two of them corresponded to large plasmids with a very similar k-mer coverage (1.69x and 1.66x) and length (107.577 kbp and 88.581 kbp). The short-read graph associated to this isolate was complex with a total of 289 nodes, 398 edges and 12 dead-ends (Figure S12).

Gplas in combination with mlplasmids predicted a total of 4 components in the plasmidome network shown in Figure S13. Two of these four components had only one connection and were formed by the small plasmids, node 84 and node 81. For these cases, gplas obtained a precision and completeness of 1.0.

The other two components corresponded to: i) a component with a precision and completeness of 1.0, formed by the nodes of the 175.881 kbp plasmid and ii) a component with a precision of 0.49 with a mixture of nodes from the 107.577 kbp and 88.581 kbp plasmid.

In the case of the last mentioned component, gplas created paths corresponding to chimeras between these two plasmids since they share repeat units and have a similar k-mer coverage that resulted in the acceptance of connections belonging to different plasmid units. For these cases, the introduction of long-read sequencing data is required to unravel the presence of two plasmid sequences present in the same predicted component.

#### REFERENCES


**1.** Antipov, D. *et al.* (2016). plasmidSPAdes : Assembling plasmids from whole genome sequencing data. *Bioinformatics*, **32**, 3380–3387.

**2.** Arredondo-Alonso, S. *et al.* (2019). Genomes of a major nosocomial pathogen enterococcus faecium are shaped by adaptive evolution of the chromosome and plasmidome. *bioRxiv*.

**3.** De Maio, N. *et al.* (2019). Comparison of long-read sequencing technologies in the hybrid assembly of complex bacterial genomes.

**4.** Decano, A. G. *et al.* (2019). Complete assembly of escherichia coli sequence type 131 genomes using long reads demonstrates antibiotic resistance gene variation within diverse plasmid and chromosomal contexts. *mSphere*, **4**(3).

**5.** Gurevich, A. *et al.* (2013). QUAST: Quality assessment tool for genome assemblies. *Bioinformatics*, **29**(8), 1072–1075.

**6.** Krueger, F. (2012). Trim galore: a wrapper tool around cutadapt and FastQC to consistently apply quality and adapter trimming to FastQ files, with some extra functionality for MspI-digested RRBS-type (reduced representation Bisufite-Seq) libraries. *URL http://www. bioinformatics. babraham. ac. uk/projects/trim_galore/. (Date of access: 28/04/2016).*

**7.** Müller, R. and Chauve, C. (2019). HyAsP, a greedy tool for plasmids identification.

**8.** Robertson, J. and Nash, J. H. E. (2018). MOB-suite: software tools for clustering, reconstruction and typing of plasmids from draft assemblies. *Microb Genom*, **4**(8).

**9.** Van Dongen, S. (2008). Graph clustering via a discrete uncoupling process. *SIAM J. Matrix Anal. Appl.*, **30**(1), 121–141.

**10.** Wick, R. R. *et al.* (2015). Bandage: Interactive visualization of de novo genome assemblies. *Bioinformatics*, **31**(20), 3350–3352.

**11.** Wick, R. R. *et al.* (2017a). Completing bacterial genome assemblies with multiplex MinION sequencing. *Microbial Genomics*, **3**(10).

**12.** Wick, R. R. *et al.* (2017b). Unicycler: Resolving bacterial genome assemblies from short and long sequencing reads. *PLoS Comput. Biol.*, **13**(6), e1005595.

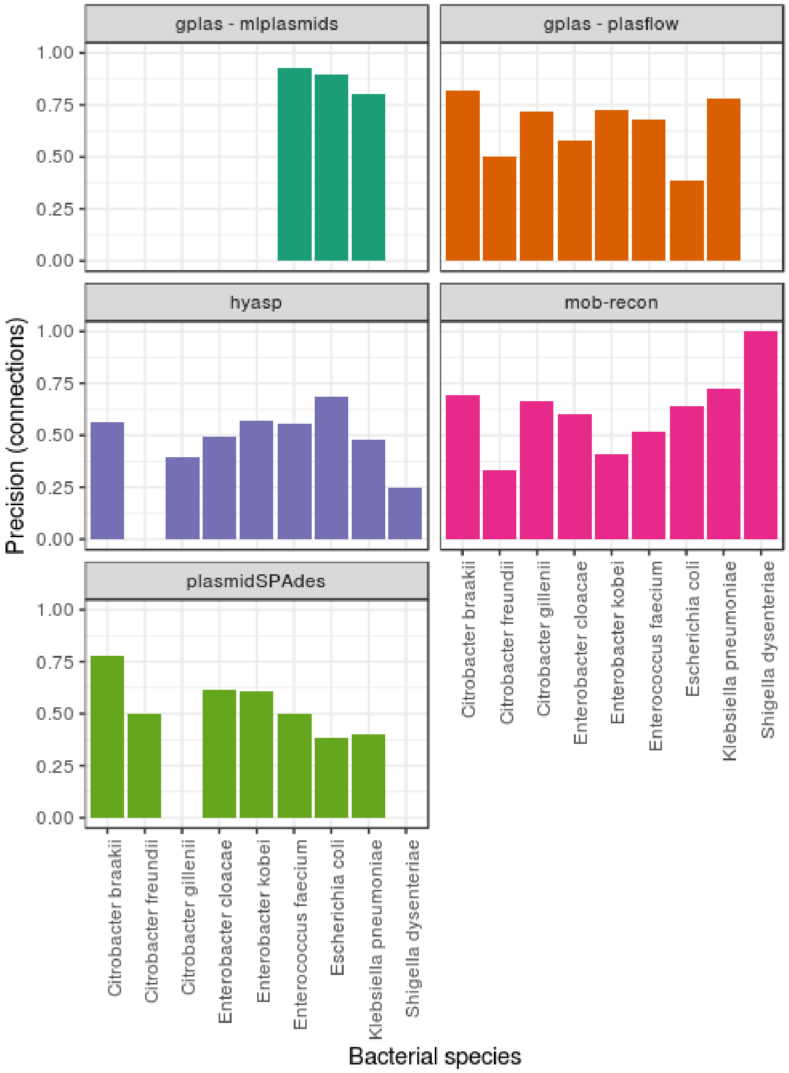


**Figure S1.** Barplot with the precision (y-axis) achieved by the different tools depending on the bacterial species analysed (x-axis).

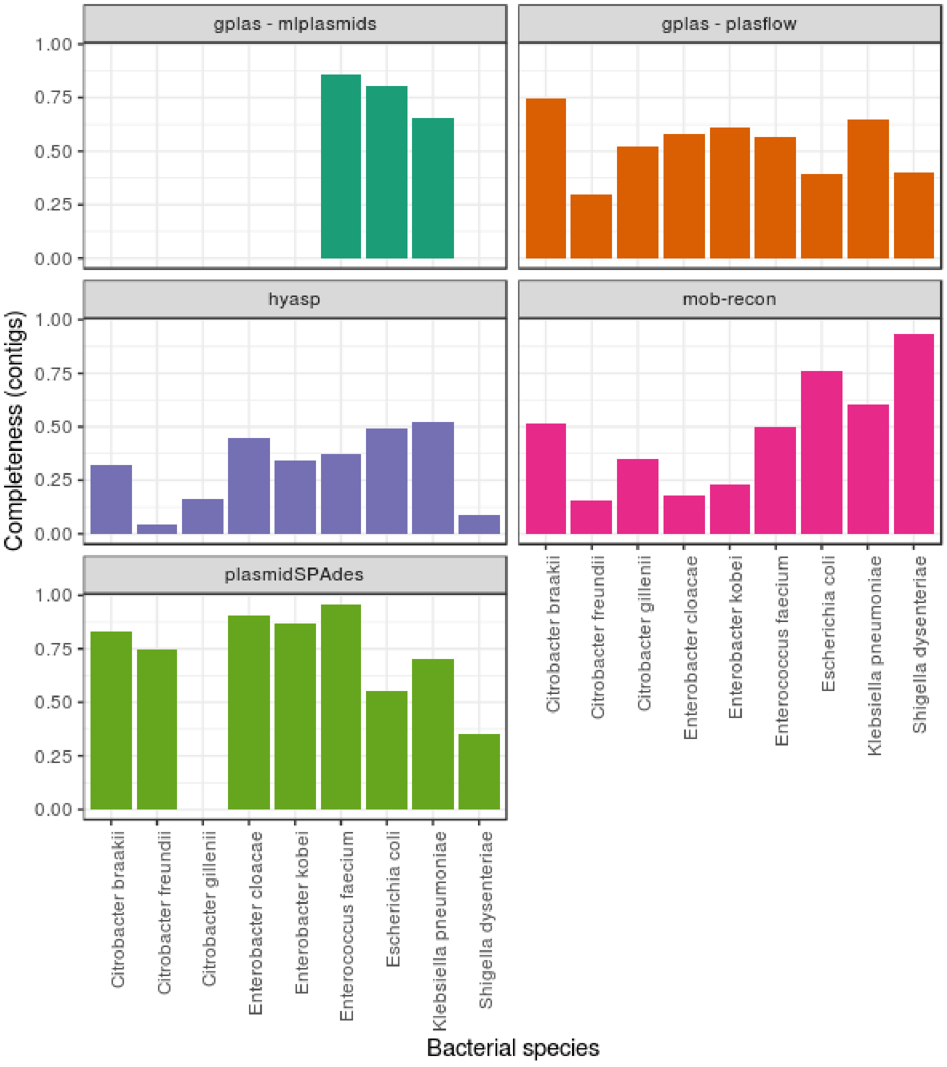


**Figure S2.** Barplot of the completeness (y-axis) achieved by the different tools depending on the bacterial species analysed (x-axis).

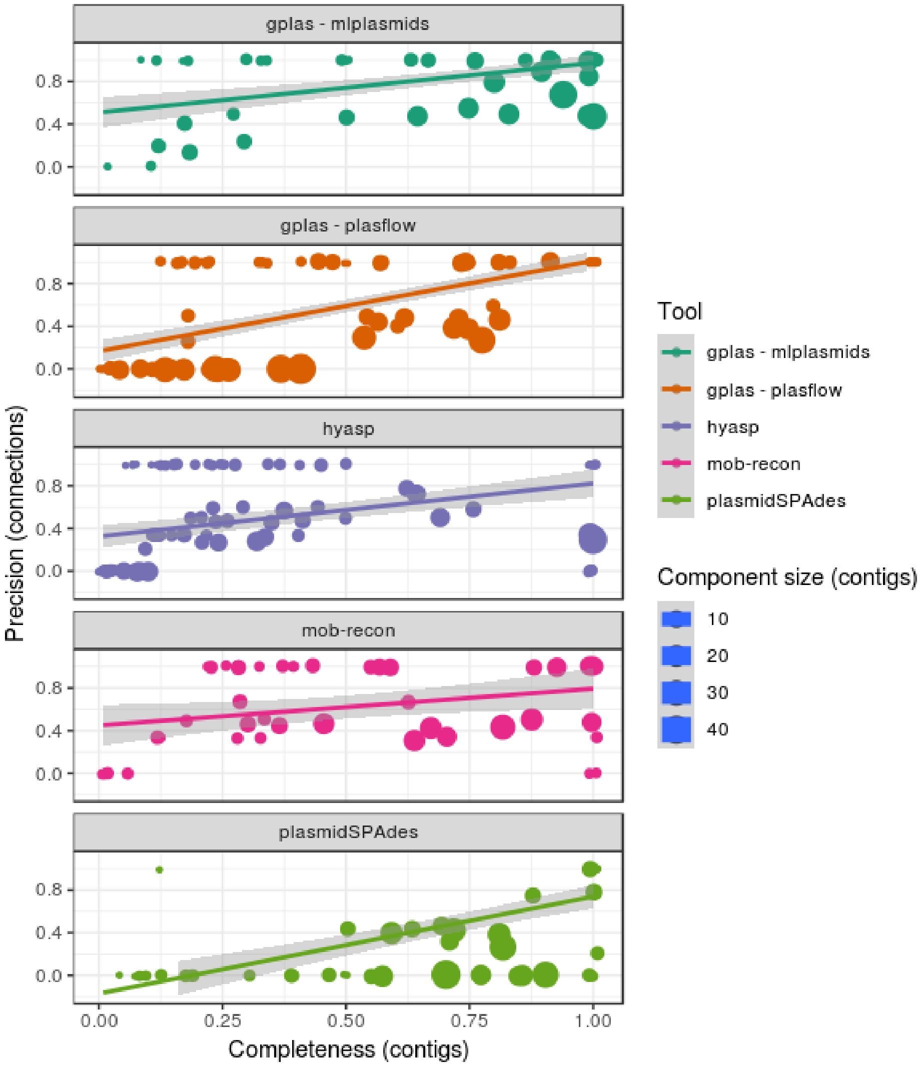


**Figure S3.** Scatterplot of the completeness (x-axis) and precision (y-axis) obtained by each component predicted by the tools included in the benchmarking. For each classifier, we fitted a linear regression model and indicated the standard error with a shadow area to observe the correlation between precision and completeness in each of the predictions.

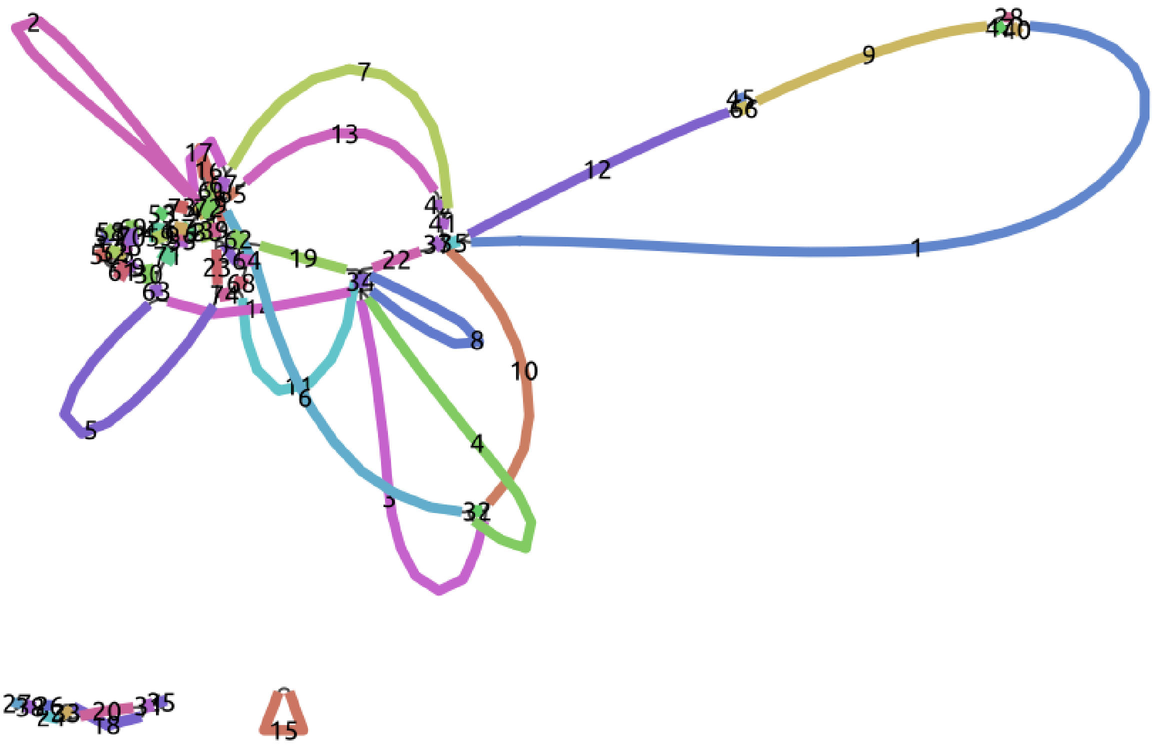


**Figure S4.** Visualization of the entire KSB1_7G isolate short-read WGS graph using Bandage.

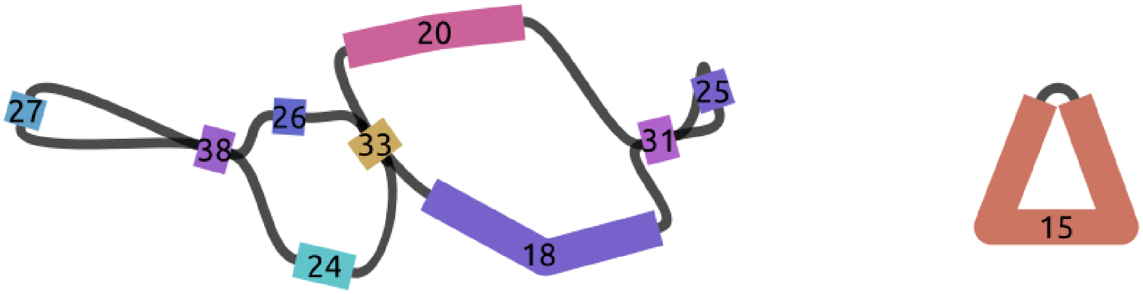


**Figure S5.** Zoom-in of the two components corresponding to plasmid sequences from KSB1_7G isolate short-read WGS graph .

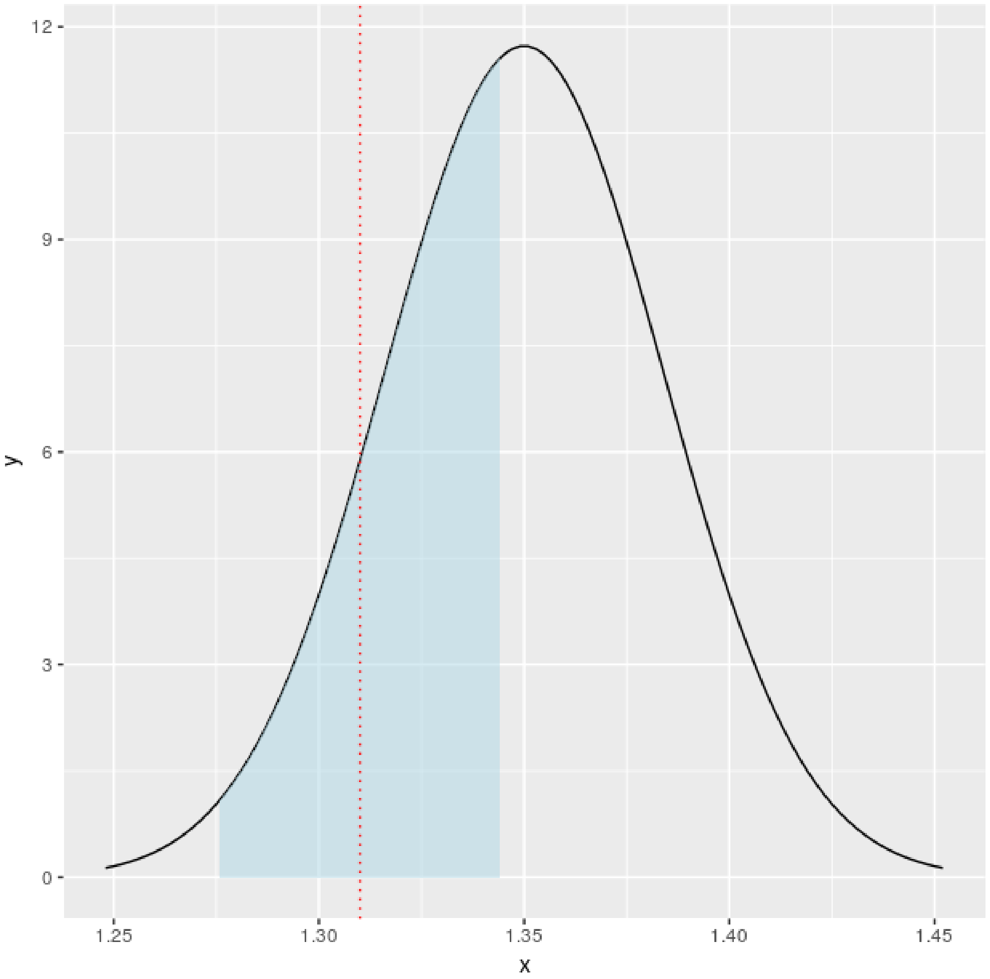


**Figure S6.** Probcoverage (0.36) of the outcoming node 20-. This score corresponds to the area under the curve between the lower limit (1.32 - 0.034) and upper limit (1.32 + 0.034) for a normal distribution with mean 1.35 and sd of 0.034. Dashed vertical red line indicates the coverage of 20-

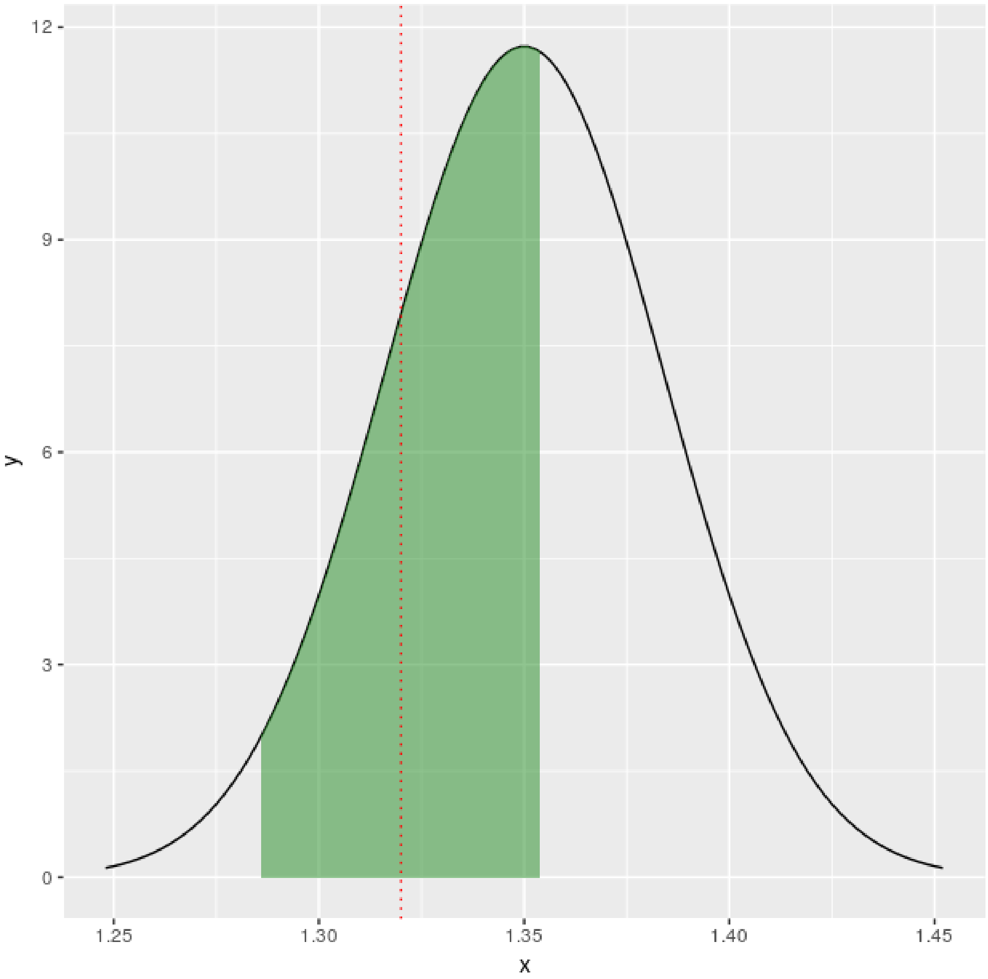


**Figure S7.** Probcoverage (0.51) of the outcoming node 24+. This score corresponds to the area under the curve between the lower limit (1.32 - 0.034) and upper limit (1.32 + 0.034) for a normal distribution with mean 1.35 and sd of 0.034. Dashed vertical red line indicates the coverage of 24+.

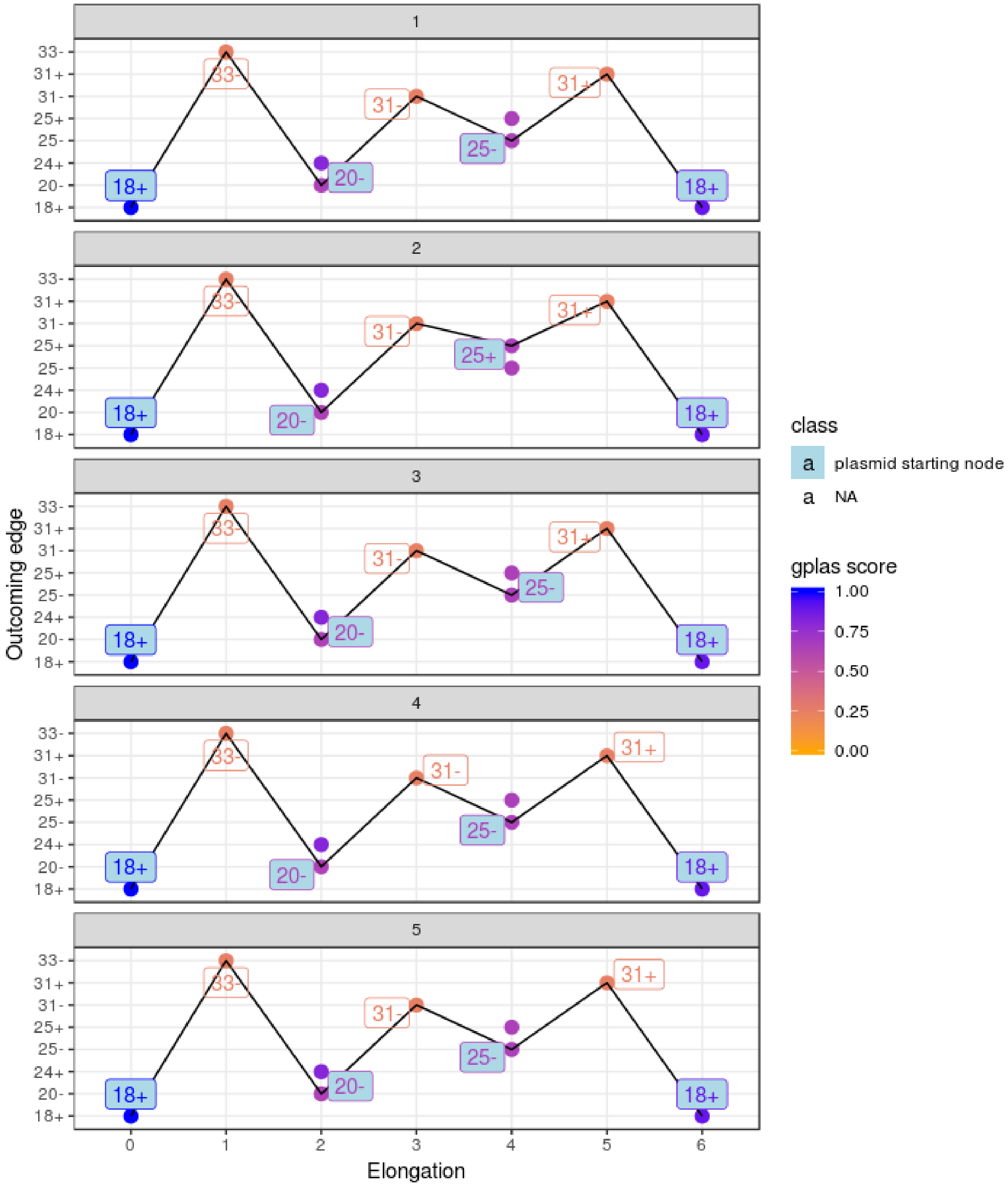


**Figure S8.** Space of solutions (n =5) starting from the plasmid starting node 18+. We observe the presence of the same solution in all the panels: 18+,33-,20-,31-,25-,31+,18+. Nodes in the solutions corresponding to plasmid starting nodes are filled with light blue.

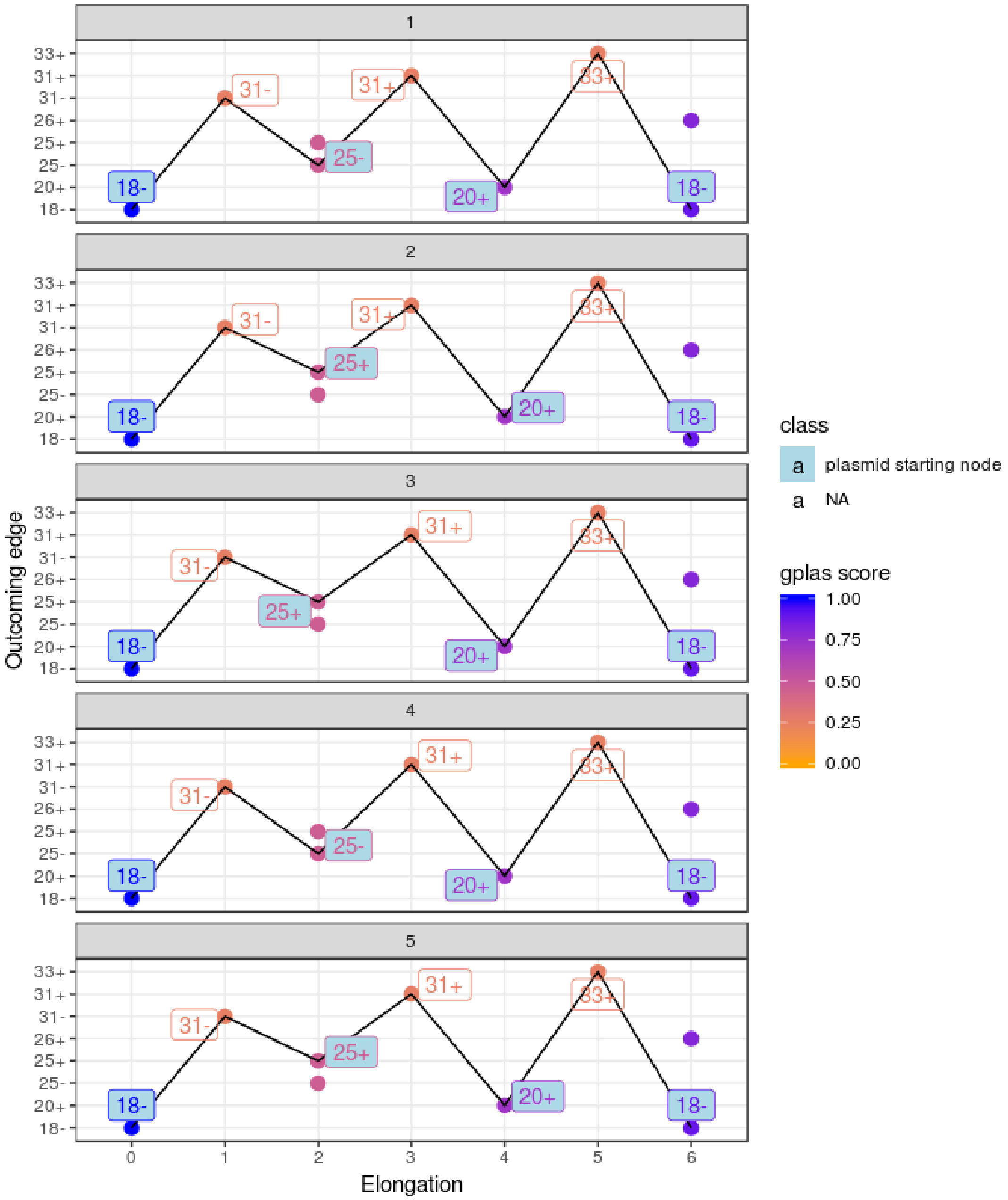


**Figure S9.** Space of solutions (n =5) starting from the plasmid starting node 18-. We observe the presence of the same solution in all the panels: 18-,31-,25+,31+,20+,33+,18-. Nodes in the solutions corresponding to plasmid starting nodes are filled with light blue.

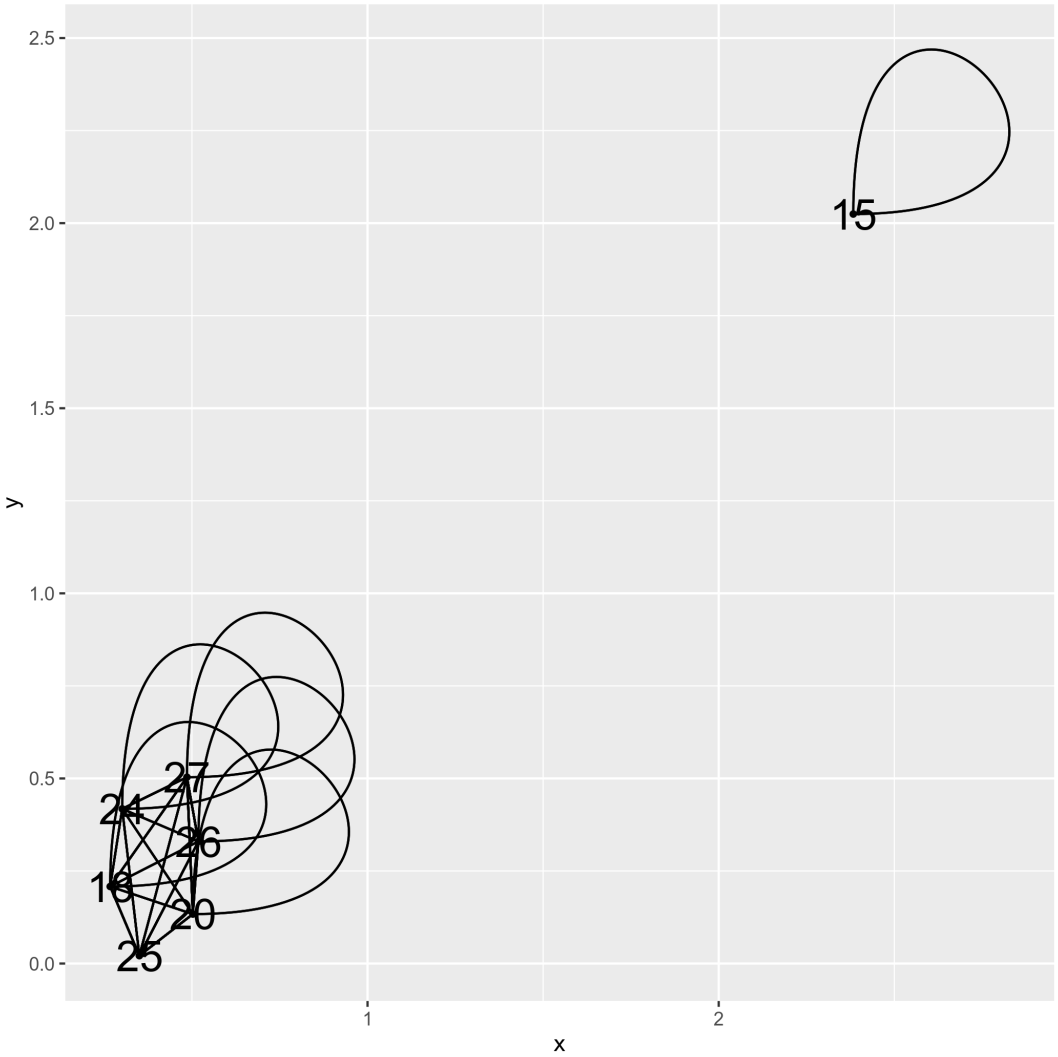


**Figure S10.** Plasmidome network from the isolate *K. pneumoniae* KSB17G. The network corresponds to an undirected graph in which nodes are the plasmid starting nodes and edges are drawn based on the co-existence of the nodes in the solutions found by gplas.

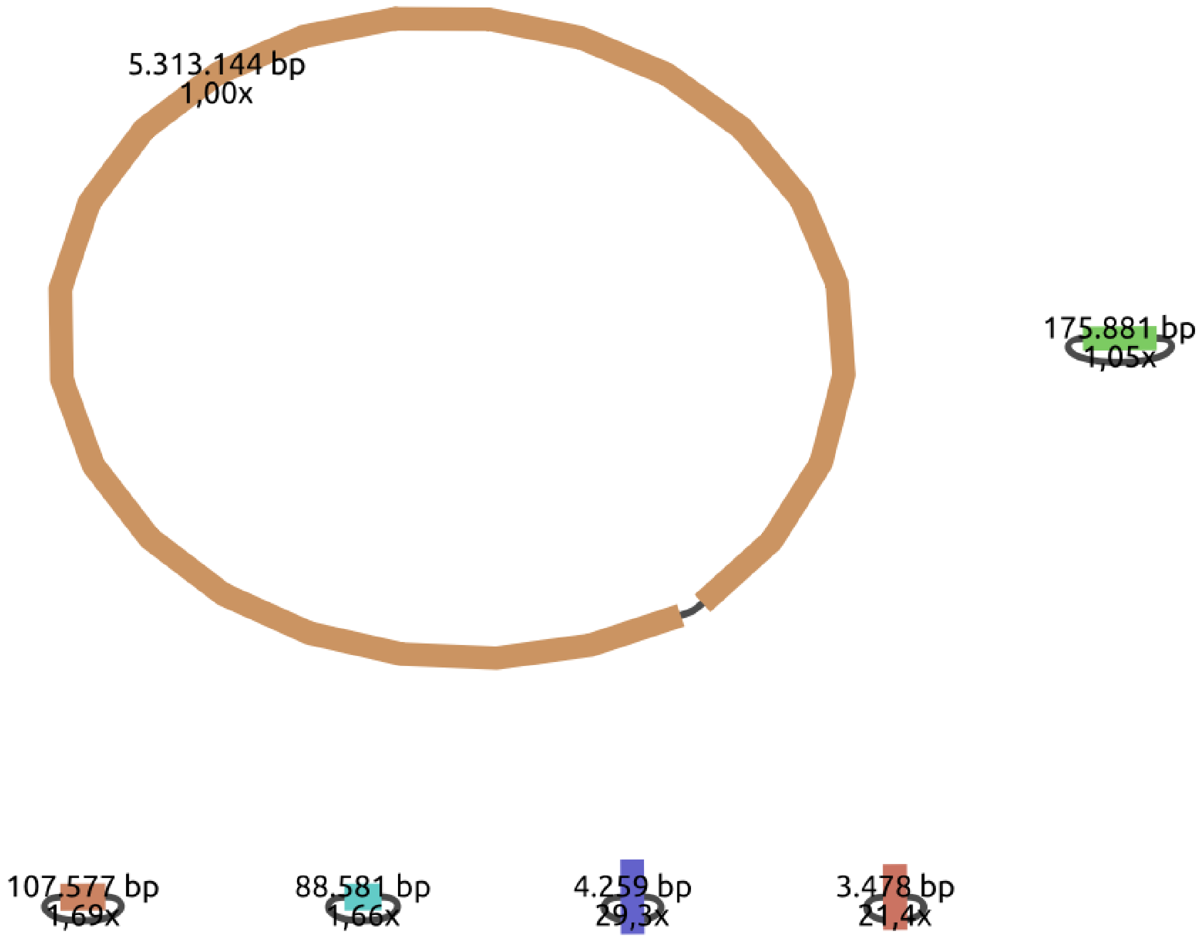


**Figure S11.** Bandage visualization of the complete genome from the *K. pneumoniae* isolate SAMN10819819 with sequence length and coverage displayed.

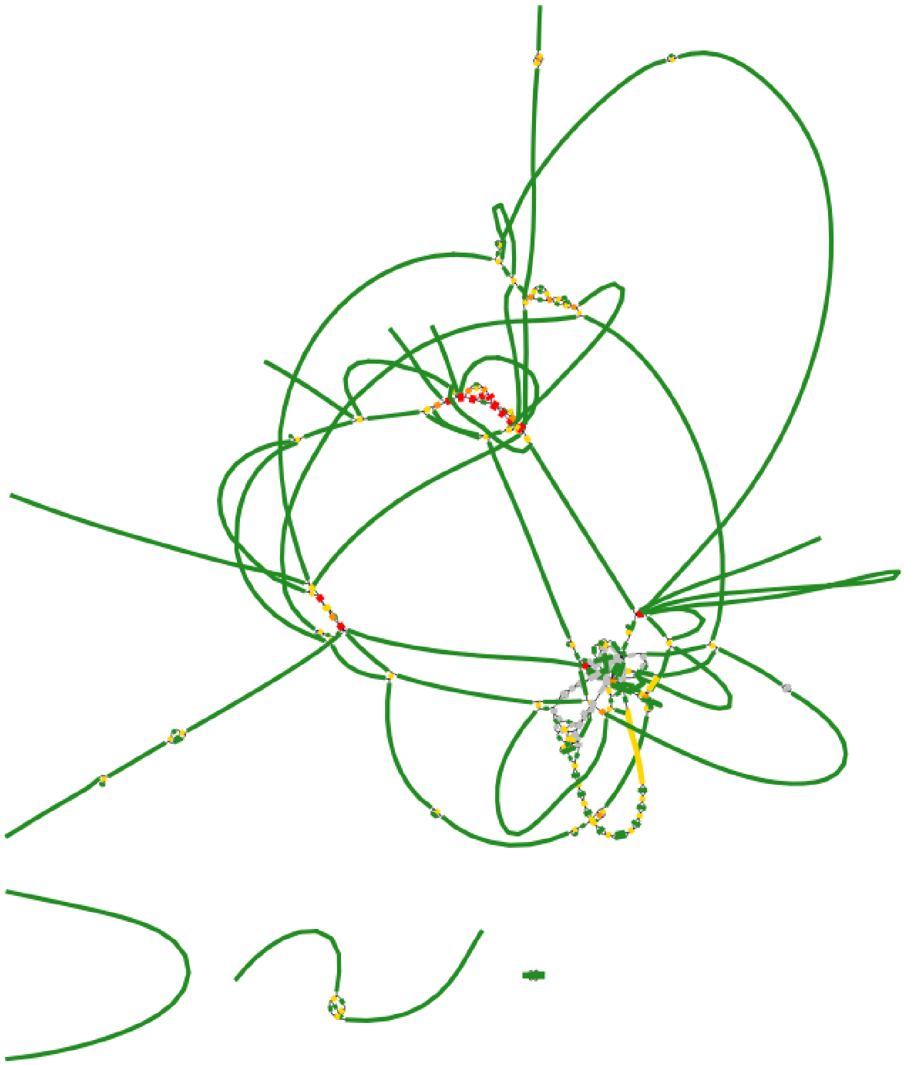

**Figure S12.** Bandage visualization of the short-read graph from the *K. pneumoniae* isolate SAMN10819819.

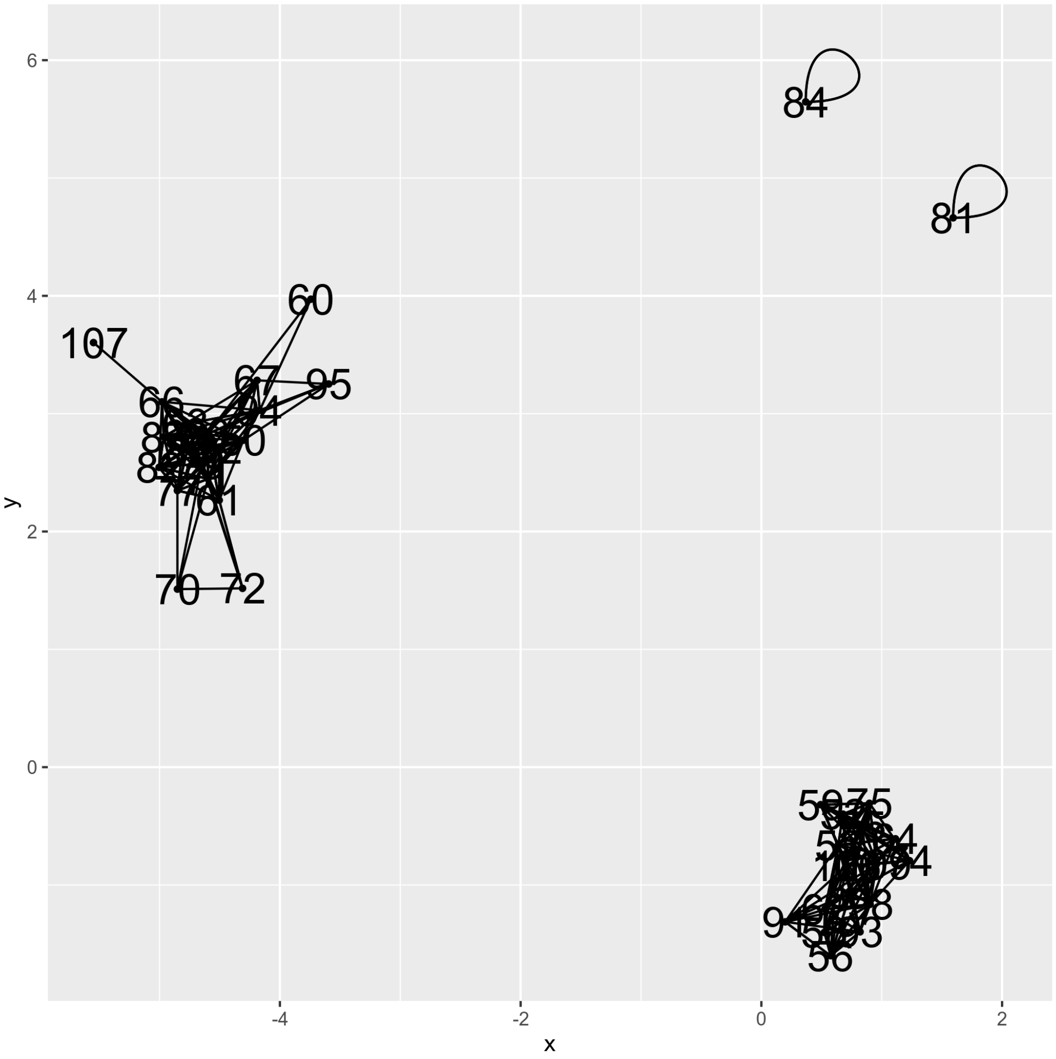


**Figure S13.** Plasmidome network created by gplas for *K. pneumoniae* SAMN10819819
